# Zinc finger domains bind low-complexity domain polymers

**DOI:** 10.1101/2023.10.29.564599

**Authors:** Naohiko Iguchi, Noriyoshi Isozumi, Yoshikazu Hattori, Tomohiro Imamura, Masatomo So, Hitoki Nanaura, Takao Kiriyama, Nobuyuki Eura, Minako Yamaoka, Mari Nakanishi, Masashi Mori, Shinya Ohki, Hiroyuki Kumeta, Hironori Koga, Mai Watabe, Takuya Mabuchi, Shingo Kanemura, Masaki Okumura, Takuya Yoshizawa, Ichiro Ota, Naoki Suzuki, Masashi Aoki, Yoshito Yamashiro, Tomohide Saio, Kazuma Sugie, Eiichiro Mori

## Abstract

Self-association of low-complexity protein sequences (LC domains) is important for polymer formation. Several molecular chaperones are involved in the regulation of LC domain polymer formation. However, the mechanisms underlying cell recognition of LC domain polymers remain unclear. Here we show that zinc finger domains (ZnFs) bind LC domains of RNA-binding proteins in a cross-β polymer-dependent manner. ZnFs bound to LC domain hydrogels and suppressed LC domain polymer formation. Moreover, ZnFs preferentially recognize LC domains in the polymeric state. These findings suggest that ZnFs act as physiological regulators of LC domain polymer formation.

## Main

Low-complexity protein sequences (LC domains) are disordered regions of proteins with a limited number of amino acids that drive phase transitions into gel-like or liquid-like states^1–3^, which in turn form membrane-less organelles and play a key role within living cells^4–6^. Dysregulating phase separation of RNA binding proteins (RBPs) with LC domains, including heterogeneous nuclear ribonucleoprotein A2 (hnRNPA2)^7^, fused in sarcoma (FUS)^8^, and TAR DNA-binding protein of 43 kDa (TDP43)^9^, leads to the formation of pathogenic fibrils and may cause amyotrophic lateral sclerosis (ALS) and other neurodegenerative diseases^10,11^.

There are several known influences on LC domains. For example, post-translational modifications control chemical properties of LC domains^12^. Karyopherins^13–16^ and other molecular chaperones^17–19^ have been found to regulate the phase separation of LC domains. A recent study also demonstrated how de novo designed proteins have the potential to inhibit the polymer formation of amyloid fibrils^20^. However, However, it remains unclear how the length of LC domain polymers are determined and subsequently kept from becoming too long.

In this study, we show zinc finger domains (ZnFs) as candidate molecules in modifying ALS pathogenesis from three bioinformatic approaches. We investigated the role of ZnFs in phase separation, with a focus on the interaction between ZnFs and LC domains. The approaches we utilized in this study include the following: i) an assessment of interactions by hydrogel binding assay; ii) monitoring the polymer formation; and iii) a detailed solution nuclear magnetic resonance (NMR) analysis. We describe here how ZnFs interact with LC domains, providing additional mechanistic insights on ZnFs.

### Genes encoding Zn-binding and ZnF are enriched in ALS with FUS-mutations

Cell-fate transition is involved in cancer, cardiovascular diseases, and neurodegenerative diseases^21,22^. To examine the molecules commonly altered, we analyzed two datasets: RNA-seq data of motor neurons (MNs) derived from *FUS* mutant human-induced pluripotent stem cells (hiPSCs)^23^ and ChIP-seq (H3K27ac) data of cell-fate transition induced by hypoxia mimic condition^24^. The genes common to both datasets included Krüppel-like factor 4 (KLF4) and other transcription factors (Fig. 1a). KLF4 is a transcription factor with ZnFs that regulates diverse cellular processes and one of the stem cell factors required for the induction of iPSCs^25^. Gene ontology (GO) enrichment analysis of two datasets revealed significant differences in the expression of genes related to transcription factors and metal binding as with genes related to cell adhesion, cytoskeleton, and cell migration. In addition, protein domain enrichment analysis in MNs derived from *FUS* mutant hiPSCs showed differences in the expression of genes with ZnFs (Extended Data Fig. 1). The genes encoding zinc binding and ZnF were enriched compared to the human proteome (Fig. 1b). These results showed that genes encoding Zn-binding and ZnF are enriched in ALS with FUS-mutations.

**Fig. 1:**
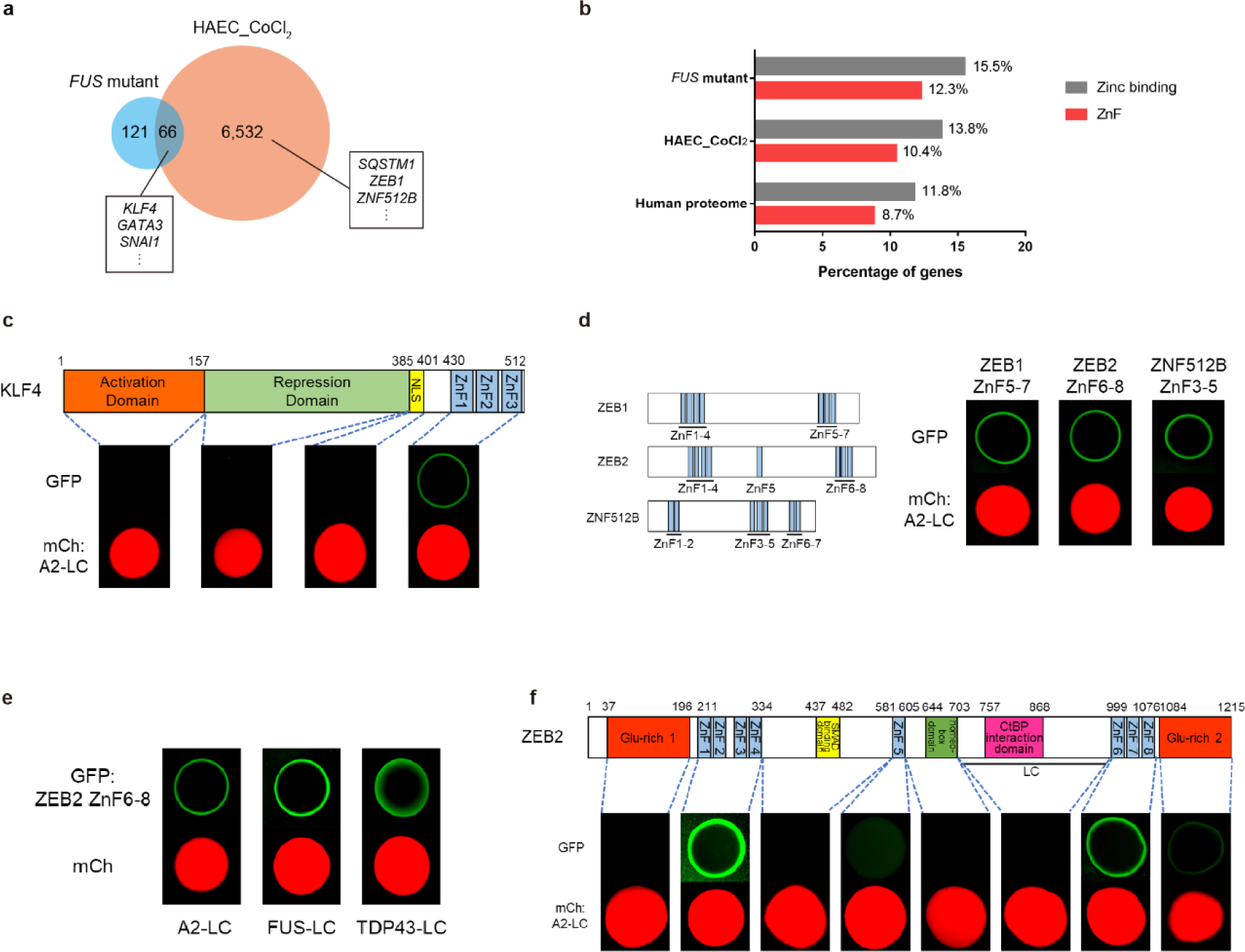
ZnFs interact with LC domains. **a.** Venn diagram showing gene overlap in two comparisons, RNA-seq data of motor neurons derived from *FUS* mutant hiPSCs (*FUS* mutant) and ChIP-seq data of CoCl_2_-treated Human aortic endothelial cell (HAEC_CoCl_2_). **b.** Percentages of genes encoding zinc binding and ZnF domains in the three datasets of FUS mutant, HAEC_CoCl_2_, and human proteome. **c**. Binding assays of GFP fusion KLF4 fragments to hydrogels derived from the mCherry fusion hnRNPA2 LC domain (mCh:A2-LC). **d**. Binding assays of various GFP fusion ZnF (GFP:ZEB1 ZnF5-7, ZEB2 ZnF6-8, and ZNF512B ZnF3-5) to mCh:A2-LC. **e**. Binding assays of GFP fusion ZEB2 ZnF6-8 (GFP:ZEB2 ZnF6-8) to various mCherry fusion LC domains (mCh:A2-LC, mCh:FUS-LC, and mCh:TDP43-LC). **f.** Binding assays of GFP fusion ZEB2 fragments to mCh:A2-LC hydrogels.

### ZnFs of transcription factors interact with LC domains of RBPs

KLF4 forms round droplets in the nucleus and has been proposed to recruit other transcription factors^26^. To test if KLF4 interacts with nuclear proteins, we performed the hydrogel binding assay. In this assay, we employed green fluorescent protein (GFP) tagged KLF4 fragments (Fig. 1c). Hydrogels of mCherry fusion LC domain of hnRNPA2 (mCh:A2-LC) were incubated with GFP-fused KLF4 fragments. Only GFP-fused ZnFs of KLF4 (GFP:KLF4-ZnF1-3) bound to the mCh:A2-LC hydrogel; other KLF4 domains did not. In addition, the DNA binding domain of other representative stem cell transcription factors, including octamer-binding transcription factor 4 (OCT4), SRY-box transcription factor 2 (SOX2) and myc proto-oncogene protein (MYC), did not bind to A2-LC (Extended Data Fig. 2a).

Zinc finger E-box binding homeobox 1 (ZEB1) and ZEB2, zinc finger protein 512B (ZNF512B) are major transcription factors and related to neurodegenerative diseases^28–29^. GFP fusion ZnFs of these transcription factors showed binding to hydrogels of mCh:A2-LC in a similar manner as seen in GFP:KLF4-ZnF1-3 (Fig. 1d). In addition, we observed that ZnFs bind to other LC domains of FUS (FUS-LC) and TDP43 (TDP43-LC) as with A2-LC (Fig. 1e). Among domains of ZEB2, ZnFs of ZEB2 bound to mCh:A2-LC hydrogels (Fig. 1f). The larger the number of ZnFs was, the stronger ZnFs bound to the A2-LC hydrogels (Extended Data Fig. 2b and 2c). ZnFs of transcription factors other than KLF4 also play an important role in binding to LC domains, and the number of ZnFs affected the recognition of LC domains.

### ZnF binds to various LC domains in a common binding manner

To investigate the interaction between ZnF and LC domains, we performed solution NMR experiments. ZnF8 of ZEB2 was used as a representative for the NMR experiments. The ^1^H-^15^N NMR spectrum of ^15^N-labeled ZnF8 showed negligible perturbations by the addition of mCherry, indicating that ZnF8 does not interact with mCherry (Fig. 2a). In contrast, the addition of mCh:A2-LC reduced the signal intensities of ZnF8, indicating that ZnF8 interacts with the LC domain of hnRNPA2 (Fig. 2a). We then assigned NMR signals of ZnF8 to identify residues of ZnF8 that interact with the LC domain of hnRNPA2 (Extended Data Fig. 3a). The addition of mCh:A2-LC eliminated ZnF8 signals, but did not change ZnF8 chemical shift values. For this reason, we used intensity ratios of ZnF8 signals to identify the binding sites of ZnF8 to mCh:A2-LC (Fig. 2b). With the addition of mCh:A2-LC, signal intensities were most significantly reduced in D1058, K1059, G1061, F1064, Q1072, H1073, and N1075.

**Fig. 2:**
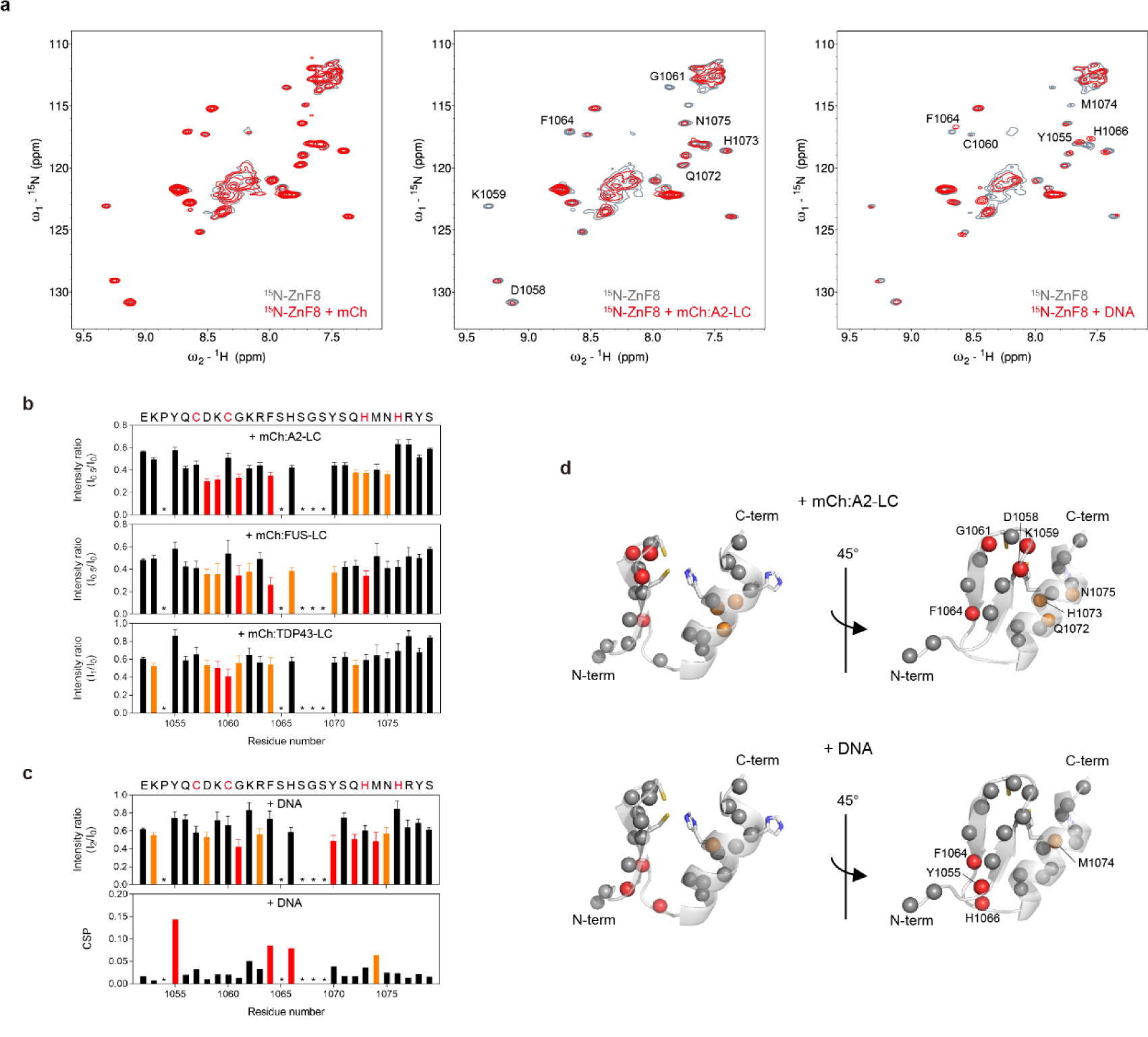
The binding mode of ZnF to LC domains is different from that to DNA. **a**. ^1^H-^15^N NMR spectra of ^15^N-labeled ZEB2-ZnF8 (^15^N-ZnF8) with mCherry (mCh; left), mCherry-fused hnRNPA2 LC domain (mCh:A2-LC; middle), ZnF-binding DNA (right). Signals labeled by amino acid type and residue number showed significant changes. **b**. Change in signal intensity of ^15^N-ZnF8 upon LC-binding. The intensity ratio is the signal intensity of LC-bound ZnF8 divided by the signal intensity of LC-free ZnF8. Significantly changed residues (intensity ratios less than mean−0.5SD and mean−SD) are indicated by orange and red bars, respectively. Residues for which no signal was detected in the absence of LC domains are indicated by asterisks. **c**. Changes in signal intensity (upper) and chemical shift (lower) of ^15^N-ZnF8 upon DNA-binding. In the graph of intensity ratios, significantly changed residues (intensity ratios less than mean−0.5SD and mean−SD) are indicated by orange and red bars, respectively. In the graph of chemical shift perturbations (CSP), significantly changed residues (CSP greater than mean+0.5SD and mean+SD) are indicated by orange and red bars, respectively. Residues for which no signal was detected in the absence of LC domains are indicated by asterisks. **d.** Ribbon model of ZEB2-ZnF8 predicted by AlphaFold2. In the upper panel, residues that were significantly changed by binding to mCh:A2-LC (intensity ratios less than mean−0.5SD and mean−SD) are shown in orange and red, respectively. In the lower panel, residues that were significantly changed by DNA-binding (CSP greater than mean+0.5SD and mean+SD) are indicated by orange and red bars, respectively.

The ZnF8 structure predicted by AlphaFold 2 has one antiparallel β-sheet and one α-helix (Fig. 2d). In this ZnF8 structure, the side chain of H1076 is not oriented to allow binding to a zinc ion. In fact, the chemical shift of H1076 side chain indicated a non-binding state with a zinc ion (Extended Data Fig. 3b). Thus, ZnF8 is likely bound to a zinc ion by three residues, C1057, C1060, and H1073, as predicted by AlphaFold 2. The interaction sites of ZnF8 with mCh:A2-LC were mapped to a loop connecting two β-strands and part of an α-helix on the predicted structure (Fig. 2d). Since hydrogel binding assays in Fig. 1e showed that ZnF can bind to various LC domains, we also analyzed the interaction of ZnF8 with LC domains of FUS and TDP43. The results showed that ZnF8 interacted with mCh:FUS-LC and mCh:TDP43-LC in a similar way with mCh:A2-LC (Fig.2b), suggesting that ZnF uses a common region for binding to various LC domains.

ZnF is known as a typical DNA-binding domain^30^. To investigate the difference between the binding of ZnF to LC domains and to DNA, the binding of ZnF8 to DNA was analyzed using NMR. The ^1^H-^15^N NMR spectrum of ZnF8 with DNA was clearly different from the ZnF8 spectrum with the LC domains (Fig. 2a). We analyzed the binding of ZnF8 to DNA using chemical shift change and revealed that Y1055, F1064, H1066, and M1074 of ZnF8 were strongly involved in DNA binding (Fig. 2c). These residues were mapped to the loop region between the β-sheet and the α-helix and part of the α-helix on the ZnF8 structure (Fig. 2d). Together, these results demonstrate that the binding region of ZnF8 to DNA is different from that to the LC domains.

### ZnFs suppressed LC domain polymer formation

We further investigated the effect of ZnF to LC domain polymer formation. First, we performed uptake experiments of GFP-fused ZnF into FUS droplets (Fig. 3a). GFP-fused ZnF8 was incorporated into the droplets more than GFP alone. We then measured the refractive index (RI) inside FUS droplets to confirm the effect of ZnF (Fig. 3b). The size of the FUS droplets was almost the same for all four conditions: no peptide, ZnF8, ZnF7-8, and ZnF6-8 (Extended Data Fig. 4). ZnF6-8 decreased RI of FUS droplets, but there was no significant difference in the RI of FUS droplets with ZnF8 and ZnF7-8 (Fig. 3c). This result suggests that the addition of ZnF6-8 makes the inside of the FUS droplets sparse. Further, we monitored the polymer formation of A2-LC using the thioflavin T (ThT), and ZnF6-8 suppressed A2-LC polymer formation (Fig. 3d).

**Fig. 3:**
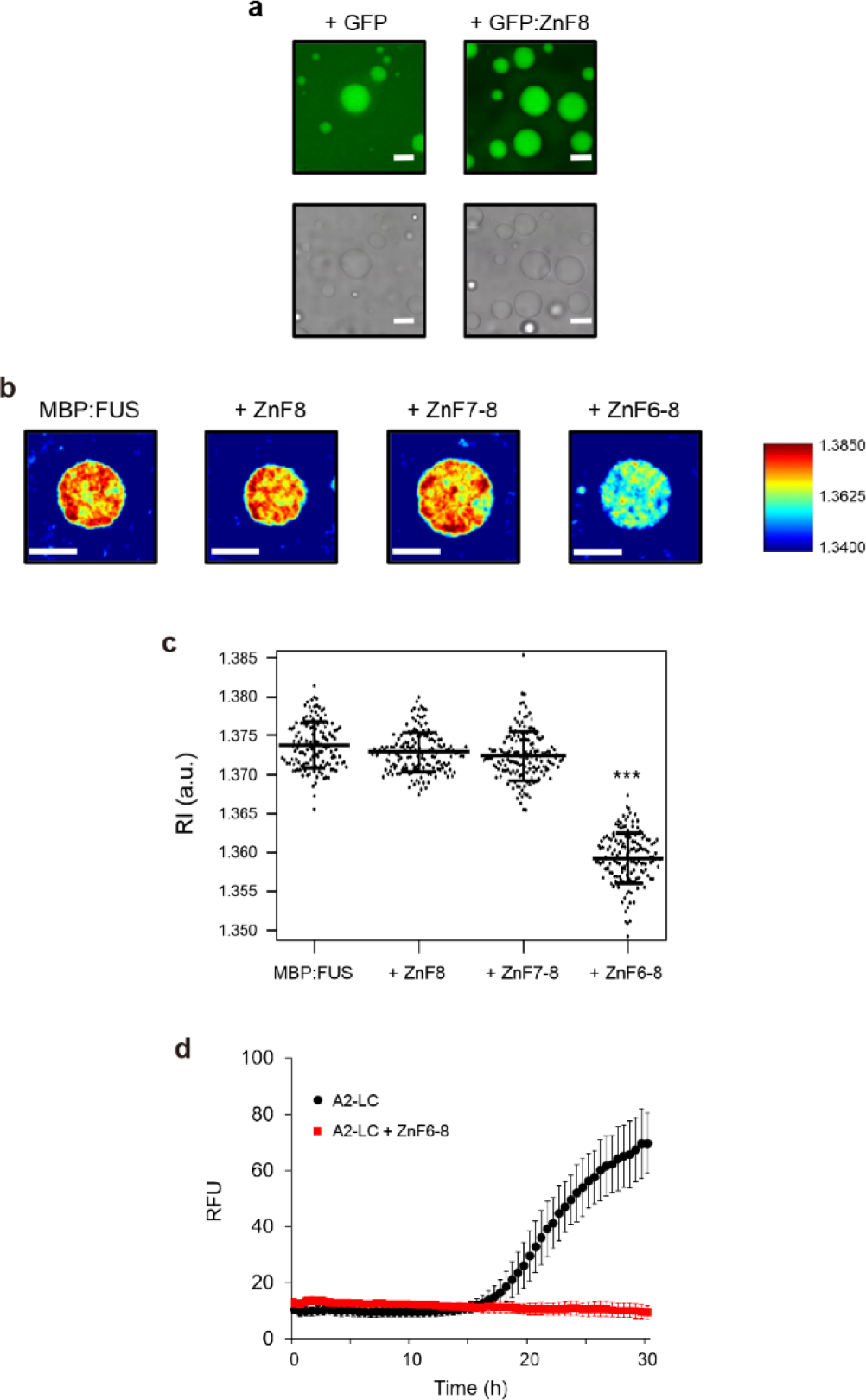
The binding of ZnF affects the phase separation of LC domains. **a.** Microscopic images of MBP:FUS droplets in the presence of GFP or GFP fusion ZEB2 ZnF8 (GFP:ZnF8). Scale bars are 5 μm. **b.** Refractive index (RI) images of MBP:FUS droplets in the absence (MBP:FUS) and presence of ZEB2 ZnF peptides (ZnF8, ZnF7-8, and ZnF6-8). Scale bars are 5 μm. **c.** Quantitative analysis of Fig. 3b. Statistical significance was examined by one-way analysis of variance with Tukey’s honest significant difference post hoc test (\*\*\**p* < 0.001). **d**. Thioflavin T assays of hnRNPA2 LC domain (A2-LC) in the absence and presence of ZEB2 ZnF6-8 peptide (ZnF6-8). Data represent means ± SD (*n* = 4).

### ZnF preferentially recognizes LC polymers over LC monomers

Our MD simulation predicted a binding model of ZnF and the LC polymers (Extended Data Fig. 5). To clarify the mechanism by which ZnF recognizes the LC domain, we tried to demonstrate whether the polymeric state of LC domain is related to ZnF binding. The F/Y to S mutation in A2-LC has been reported to suppress polymer formation^31^. In addition, we prepared a A2-LC mutant (A2-LC_MM) in which multiple F/Y residues were mutated to S (Y257S, Y264S, Y278S, F291S, F309S, and Y319S). The interaction of these mutants with ZnF8 was monitored by NMR spectra (Fig. 4a). With the addition of mCh:A2-LC and mCh:A2-LC_MM, K1059 and G1061 showed significant changes in ZnF8 signals, while Y1070 showed little change (Fig. 4b). Overall, the intensity ratio of ZnF8 signals was smaller in the order of mCh:A2-LC, mCh:A2-LC_F291S, and mCh:A2-LC_MM (Fig. 4b). These results suggest that ZnF preferentially recognize LC domains in the polymeric state. We offer a model that ZnFs act as physiological regulators of LC domain polymer formation (Fig. 4c)

**Fig. 4:**
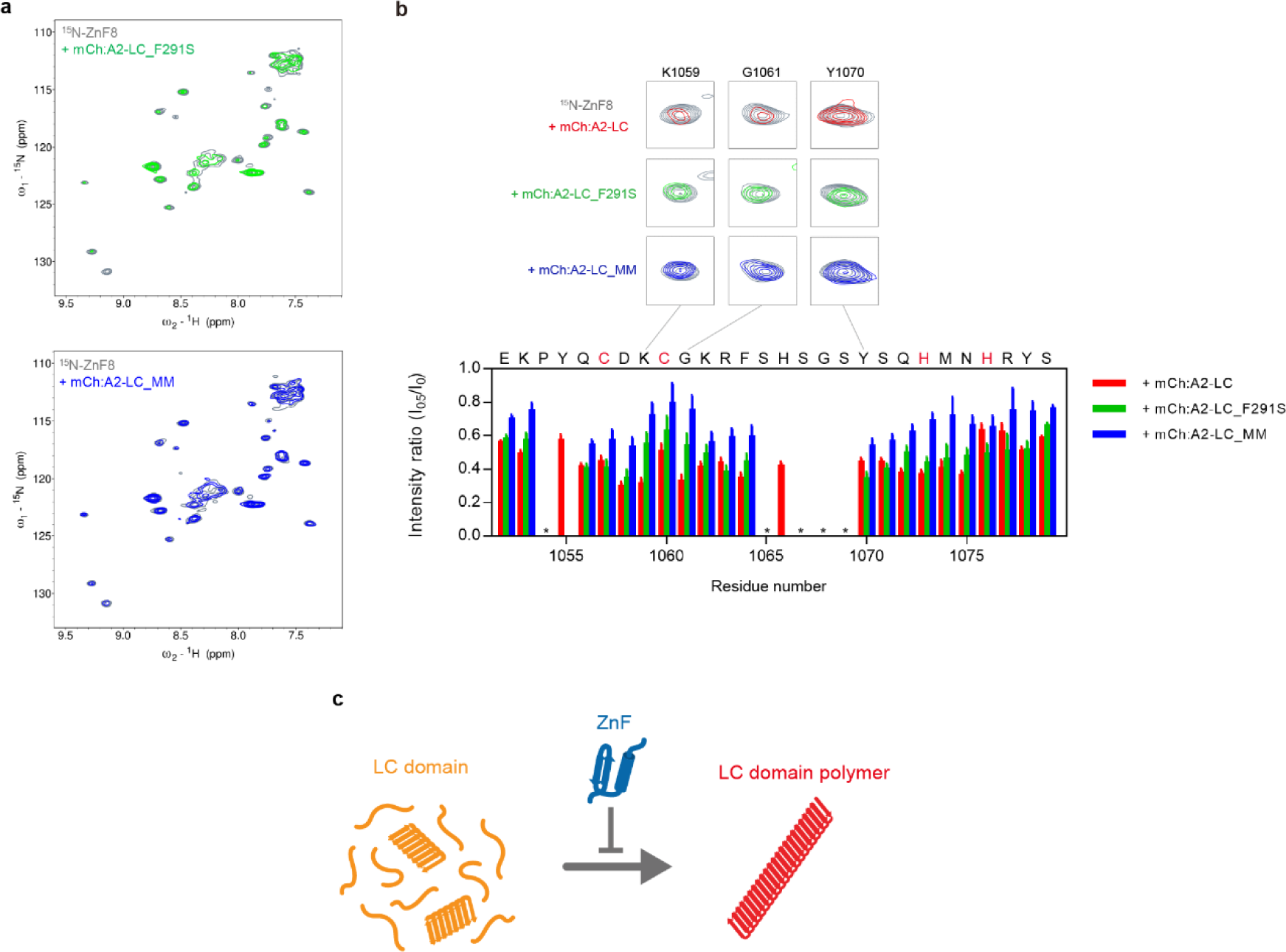
ZnF preferentially recognizes LC polymers over monomeric LC domains. **a**. ^1^H-^15^N NMR spectra of ^15^N-labeled ZEB2 ZnF8 (^15^N-ZnF8) with mCherry fusion hnRNPA2 LC domain mutants (mCh:A2-LC_F291S and mCh:A2-LC_MM). **b.** Change in signal intensity of ^15^N-ZnF8 in the addition of mCh:A2-LC (red), mCh:A2-LC_F291S (green), and mCh:A2-LC_MM (blue). The intensity ratio is the signal intensity of A2-LC-bound ZnF8 divided by the signal intensity of A2-LC-free ZnF8. Residues for which no signal was detected in the absence of LC domains are indicated by asterisks. In the upper panel, the NMR signals corresponding to the three residues K1059, G1061, and Y1070 are shown. **c.** a model that ZnFs play as physiological regulators of LC domain polymer formation.

## Discussion

ZnF is a widespread domain structure. ZnFs with two cysteins and two histidines (C2H2-type) are mostly found in transcription factors and recognize DNA^32^. Some C2H2-type ZnFs interact with other proteins and play an important role in their communication with other proteins that are attracted to the multimolecule transcriptional complex formed on DNA^33^. Through our experiments, we found that ZnFs preferentially recognize LC domains in the polymeric state, and the interaction sites of ZnF with LC domains were mapped to a loop connecting two β-strands. De novo designed proteins binding amyloid fibrils contain one α-helix and two or three β-strands, similar to the structure of C2H2-type ZnF, and the primary interaction between these designed proteins and amyloid fibrils is the stacking of a β-strand of the designed proteins onto the β-strand of the amyloid fibrils^20^. ZnF structure with one antiparallel β-sheet and one α-helix surrounding zinc ion may be important in the identification of LC domain polymers.

We found that FUS droplets absorbed ZnF and began to disappear, and that the polymer formation of LC domains was suppressed in the presence of ZnFs in ThT assay. Molecular chaperones, including karyopherin-β2^13,15^, cyclophilin A^31^, and heat shock proteins^17–19^ interact with LC domains or other segments of RBPs and regulate phase separation. ZnFs are new factors in phase separation with the specific function recognizing LC domain polymers and regulating phase separation suppressively.

Transcription factors with ZnFs, such as KLF4, ZEB1 and ZEB2, not only regulate the specific expression of human genome in the brain^34^ but also are involved in neural development and neurodegenerative diseases^35–38^. Proteins with ZnFs or zinc binding lesions have been suggested to be involved in phase separation^39–41^. Further elucidation of the function of ZnFs, including interaction with LC domains, may lead to an understanding of the pathology of neurodegenerative diseases and the invention of therapeutic methods.

## Methods

### Bioinformatic analysis

RNA-seq of MNs derived from *FUS^H^*^517^*^D/H^*^517^*^D^* mutant hiPSCs was used for bioinformatic analyses^23^. Among the differential expressed genes (DEGs) in MNs (SDs and Axon), those with a statistically significant difference (*P* < 0.1) were selected. Our previous ChIP-seq (H3K27ac) data (GSE189071) of cobalt chroride (CoCl_2_)-treated (hypoxia mimic) human aortic endothelial cells (HAEC) was also used for bioinformatic analyses^24^. RNAs that did not express genes such as antisense RNAs, microRNAs, non-cording RNAs were removed from all data. For gene ontology (GO) analysis and domain enrichment analysis (InterPro), the DAVID server was used. Due to the limited number of input genes on the DAVID server, the ChIP-seq data used the top 6,000 genes in the score. For zinc binding and ZnF enrichment analysis, proteins with zinc binding or ZnF domain were annotated using the UniProt database (keyword: KW-0862 and KW-0863). The human proteome dataset for gene counts was obtained from UniProt sever (Proteome ID: UP000005640; release 2022_04).

### Expression and purification of recombinant proteins

Expression plasmids for His-mCherry fusion LC domains of hnRNPA2 (residues 181– 341, mCh:hnRNPA2-LC) and FUS (residues 2–214, mCh:FUS-LC) were obtained from the Steven L. McKnight Laboratory. Expression plasmids for other His-mCherry or His-GFP fusion proteins were constructed using in-Fusion HD Cloning Kit (Takara Bio, Inc.). All mCherry/GFP fusion proteins were expressed in *E. coli* BL21(DE3) cells with 0.5 mM isopropyl-β-D-thiogalactopyranoside (IPTG). MCh:hnRNPA2-LC (residues 181– 341), hnRNPA2-LC mutants (mCh:hnRNPA2-LC_F291S and mCh:hnRNPA2-LC_MM), and mCh:FUS-LC (residues 2–214) were expressed at 20°C overnight, and mCh:TDP43-LC (residues 262–414) was expressed at 16°C overnight. All mCherry fusion proteins were purified by Ni-NTA agarose (FUJIFILM Wako Pure Chemical Corporation) according to steps described in our previous study^42^. GFP:KLF4-AD (residues 1-157), GFP:KLF4-RD (residues 158-385), GFP:KLF4-NLS (residues 430-512), GFP:KLF4-ZnF1-3 (residues 430-512), GFP:SOX2-HMG (residues 41-109), GFP:MYC-bHLH-LZ (residues 368-448), GFP:OCT4-POU (residues 138-289), GFP:ZEB1-ZnF5-7 (residues 904-981), GFP:ZEB2-ZnF6-8 (residues 999-1076), GFP:ZNF512B-ZnF345 (residues 540-653), GFP:OCT4-POU (residues 138-289), GFP:ZEB2-G-rich1 (residues 37-196), GFP:ZEB2-ZnF1-4 (residues 211-334), GFP:ZEB2-SMAD (residues 335-580), GFP:ZEB2-ZnF5 (residues 581-605), GFP:ZEB2-homeobox (residues 606-703), GFP:ZEB2-LC (residues 704-998), GFP:ZEB2-G-rich2 (residues 1084-1215), GFP:ZEB2-ZnF6-7 (residues 999-1049), GFP:ZEB2-ZnF7-8 (residues 1027-1076), GFP:ZEB2-ZnF6 (residues 999-1021), GFP:ZEB2-ZnF7 (residues 1027-1049), and GFP:ZEB2-ZnF8 (residues 1055-1076) were expressed at 20°C overnight and purified using the same method as mCherry fusion proteins. Briefly, cells were lysed in a buffer containing 25mM Tris-HCl pH 7.5, 200mM NaCl, 10mM β-ME, and Protease Inhibitor Cocktail Tablets. GFP fusion proteins were purified using Ni-NTA agarose with a wash buffer (25mM Tris-HCl pH 7.5, 200mM NaCl, 10mM β-ME, and 20mM imidazole) and elution buffer (25mM Tris-HCl pH 7.5, 200mM NaCl, 10mM β-ME, and 300 mM imidazole). Where necessary, 2 M Urea was added to the buffers.

For His-hnRNPA2, the expression plasmid was constructed using in-Fusion HD Cloning Kit with pHis-parallel1 vector as the template. His-hnRNPA2-LC was expressed in *E. coli* BL21(DE3) cells with 0.5 mM IPTG at 23°C overnight. Cells were lysed in buffer (50 mM Tris-HCl pH 7.5, 500 mM NaCl, 10mM β-ME, Protease Inhibitor) and centrifuged at 12,000×g for 30 minutes. After centrifugation, the pellet was collected and dissolved in solubilization buffer (50 mM Tris-HCl pH 7.5, 500 mM NaCl, 10mM β-ME, and 6 M guanidine) followed by Ni-NTA purification.

### Hydrogel binding assays

Hydrogel droplets of mCh:hnRNPA2-LC, mCh:FUS-LC and mCh:TDP43-LC were prepared according to procedures reported in a previous study^43^. For hydrogel binding assays, purified GFP fusion proteins were diluted to 1 μM in the buffer (20mM Tris-HCl pH 7.5, 150mM NaCl, 20mM β-ME, 0.1mM phenylmethylsulfonyl fluoride (PMSF), and 0.5mM EDTA) and pipetted onto a hydrogel dish. Horizontal sections of the droplets were scanned with excitation wavelengths on a confocal microscope (FLUOVIEW FV3000, OLYMPUS) after overnight incubation. We used the profile plot mode in ImageJ and measured relative intensity of GFP signals across the hydrogel droplets in triplicate. The values are presented as mean ± SD. Graphpad Prism version 7 was used for statistical analysis.

### ZnF peptides

The GST:ZnF expression plasmid was transformed into *E. coli* BL21(DE3) cells. *E. coli* cells with the plasmid were cultured in LB/Kan medium at 37°C, and protein expression was introduced by the addition of 0.5 mM IPTG. After 3 h of incubation at 37°C, cells were collected and suspended in PBS containing 0.1 mM EDTA and protease inhibitor. For ^15^N-labeling or ^13^C,^15^N-double labeling, *E. coli* cells were respectively cultured in M9/Kan medium with ^15^NH_4_Cl, or ^15^NH_4_Cl and ^13^C-glucose at 37°C. The ^15^N-labeled protein was expressed by the addition of 0.5 mM IPTG followed by a 4 h incubation at 37°C. Cells were sonicated and centrifugated at 12,000×g for 30 min at 4°C. GST-fused proteins were purified using Glutathione Sepharose 4B beads (Cytiva) and GST-tag was cleaved with PreScission Protease (Cytiva) at 4°C for 16 h. ZnF peptides were purified using gel filtration (Superdex 30 pg16/600, Cytiva) with a buffer (20 mM Tris-HCl pH 8.0, 200 mM NaCl, and 1 mM DTT), and desalted using Sep-Pak C18 (Waters). The desalted peptides containing acetonitrile were dried under vacuum conditions.

### ZnF-binding DNA

Two complementary single strand oligonucleotides (5’-GTAATCTGGGCCACCTGCCTGGGAGGA-3’ and 5’-TCCTCCCAGGCAGGTGGCCCAGATTAC-3’) corresponding E2 box_44_ were Two complementary single strand oligonucleotides (5’-purchased from Sigma-Aldrich. An equal molar ratio of single strand oligonucleotides was mixed and incubated at 95°C for 5 min. The double-stranded DNA obtained by this annealing was used for NMR experiments.

### NMR experiments

For titration experiments, the ^15^N-labeled sample at 0.1 mM concentration were dissolved in a solution containing 20 mM Tris-HCl pH 7.5, 200 mM NaCl, 20 mM β-ME, 0.1 mM PMSF, 0.12 mM ZnCl_2_, and 5% ^2^H_2_O. NMR spectra were measured on the Bruker AVANCE 500 MHz spectrometer. The sample temperature was set at 15°C. ^15^N-SOFAST HMQC pulse sequence was employed to obtain backbone ^1^H-^15^N correlation spectrum. The following samples were used for the titration experiments; mCherry, mCherry:A2-LC, mCherry:FUS-LC, mCherry:TDP43-LC, and ZnF-binding DNA. The proteins and DNA were prepared at 1 mM and titrated into ZnF8 at the molar ratios of 0.2, 0.5, 1, and 2.

For backbone resonance assignment experiments, the ^13^C,^15^N-labeled sample at 0.9 mM concentration were dissolved in a solution containing 20 mM Tris-HCl pH 7.5, 20 mM β-ME, 0.1 mM PMSF, 1 mM ZnCl_2_, and 5% ^2^H_2_O. NMR spectra were measured on Bruker AVANCE III 600 MHz equipped with a TBI probe and AVANCE NEO 800 MHz spectrometers equipped with a CPTCI probe. The sample temperature was set at 10°C. Backbone resonance assignment was carried out using the following triple resonance spectra: HNCO, HNCA, HN(CO)CA, HNCACB, and CBCA(CO)NH. The assignment was transferred to titration experiment conditions by tracing the change of sample concentration. Data were processed using NMRPipe^45^, and the spectra were analyzed using NMRFAM-Sparky^46^ and MagRO-NMRView^47^. The standard deviation (*SD*) of intensity ratio was estimated from the signal to noise ratio (*SNR*) using the following equation:

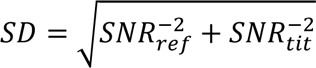

where 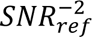 and 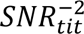 are the signal to noise ratios of the peak on the reference spectrum and titrated spectrum, respectively. The chemical shift perturbation (*CSP*) of the backbone amide group was averaged using the following equation:

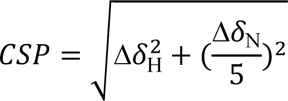

where Δδ_H_ and Δδ_N_ are the chemical shift changes of proton and nitrogen, respectively.

### Refractive index measurement inside FUS droplet using the holotomography microscope

The LLPS-droplet formation was initiated by mixing 10 μM of MBP:FUS with 8% polyethylene glycol 8000 in a buffer containing 20 mM Tris-HCl, pH 7.5, 200 mM NaCl, 10% glycerol, and 20 mM β-ME. After 1 min, 10 μM of ZnF8, ZnF7-8, or ZnF6-8 was added to the mixture and incubated for 5 min. For the GFP-fluorescence imaging, 10 μM of GFP:ZnF8, GFP:ZnF7-8, or GFP:ZnF6-8 was added to the mixture and incubated for 5 min. The droplets were observed using a holotomography microscope with laser-induced fluorescence system (Tomocube, Inc.)^48^. The refractive index (RI) and fluorescence image show the XY cross section of the droplet center. Average RI inside the droplet and their radius were calculated by enclosing the RI image in a circle using TomoStudio version 3.2.8 software (Tomocube, Inc.) and plotted by the Igor Pro version 6.36 software (WaveMetrics). The statistical significance of differences was examined by a one-way analysis of variance with Tukey’s honest significant difference (HSD) post hoc testing. All statistical tests were performed using KaleidaGraph version 4.5.1 software (Synergy Software) at a significance level of α = 0.05.

### ThT fluorescence assays

His-tagged hnRNPA2-LC was dissolved in stock buffer containing 20 mM sodium phosphate, pH 7.4, 500 mM NaCl, and 8 M Urea. The sample was diluted to 20 μM with an assay buffer containing 20 mM Tris-HCl, pH 7.5, 200 mM NaCl, 20 mM β-ME, 0.1 mM PMSF, 1 mM ZnCl_2_, and 20 μM ThT. In the condition with ZnF, a sample containing 20 μM ZnF6-8 was prepared. ThT fluorescence signal was monitored with a Varioskan LUX plate reader (Thermo Fisher Scientific) using excitation and emission wavelengths of 440 and 484 nm, respectively. Fluorescence data were analyzed using Skanlt software (Thermo Fisher Scientific).

### MD simulation

Molecular docking simulations were performed using the AutoDock Vina 1.1.2 software^49^ on the ZnF8 and hnRNPA2 complex. The ZnF8 domain between residues Glu1052 and Ser1079 was extracted from the molecular structures of the ZEB2 estimated by the AlphaFold 2^50^. The cryo-electron microscopy structure of the low-complexity (LC) domain of hnRNPA2 (PDB ID: 6WQK)^51^ was used. The size of the grid boxes was chosen based on its ability to occupy the whole ZnF8 and hnRNPA2 complex. The molecular dynamics (MD) simulations of ZnF8 bound to the LC domain of hnRNPA2 were then performed with the CHARMM36m force field^52^ using LAMMPS software^53^. The selected docked poses provided in the docking simulations were used as starting structures, placed in a periodic simulation box and solvated with water. The salt concentration was set to 150 mM NaCl, and corresponding numbers of Na+ and Cl− ions were added. After the steepest-descent energy minimization, production runs were performed for 100 ns in the NPT ensemble at 300 K and 1 atm, and trajectory data were collected every 10 ps. During these MD simulations, C’ atoms of terminal residues and Cα atoms at the beginning and end of β sheet were fixed to maintain the secondary structures of the LC domain of hnRNPA2. The particle−particle particle−mesh (PPPM) method^54^ was used to calculate long-range electrostatic interactions. Equations of motion were integrated using the Verlet algorithm^55^ with a time step of 2 fs, along with the SHAKE algorithm^56^ to constrain the lengths of bonds to hydrogen. The binding free energy between ZnF8 and hnRNPA2 was calculated by the MM-GBSA method^57^ using the MD simulation trajectories over the last 50 ns. A model that has the lowest binding free energy among the estimated binding sites is used as the most plausible binding model.

## Acknowledgements

This work was supported by grants from AMED [JP23wm0425004 to E.M.; JP23ek0109642 to T.S.; JP23ek0109558 to T.Y. and H.N.; JP21ek0210158 to Y.Y.], JSPS KAKENHI [JP20H03199 to E.M.; JP19K17044 to N. Iguchi; JP20KK0156, JP22K18361, and JP22H02560 to T.S.; JP23K05657 to Y.H.; JP22K15734 to H.N.; JP21H02677 to Y.Y.; JP22K15691 to M.N.; JP21H05095, JP22H02205, JP23KK0105 to M.O.], MEXT Grant-in-Aid for Transformative Research Areas (B) [JP21H05094 and JP21H05093 to T.S.], JST FOREST Program [JPMJFR204W to T.S.; JPMJFR201F to M.O.]; Takeda Science Foundation to E.M. and T.S.; Kato Memorial Bioscience Foundation to H.N.; Uehara Memorial Foundation to M.O. and S.K. This work was partially supported by Joint Usage and Joint Research Programs, Institute of Advanced Medical Sciences, Tokushima University. This work was supported in part by “Advanced Research Infrastructure for Materials and Nanotechnology in Japan (ARIM)” of the Ministry of Education, Culture, Sports, Science and Technology (MEXT) under Grant Number JPMXP12-22JI0044. We thank Shunsuke Tomita (AIST), Shinsuke Niwa (Tohoku University), Yoichi Shinkai (AIST), Wataru Iwasaki (Tokyo University), and Noriyuki Kodera (Kanazawa University) for technical support. NMR experiments were performed at the Hokkaido University Advanced NMR Facility, a member of the NMR Platform.

## Author contributions

N. Iguchi, N. Isozumi, K. Sugie, and E.M. designed the research. N. Iguchi, N. Isozumi, Y.H., T.I., M.S., H.N., M.M., H. Kumeta., H. Koga, M.W., T.M., S.K., M.O., T.Y., Y.Y. and T. Saio performed research. N. Iguchi, N. Isozumi, Y.H., T.M., M.S., M.W., S.K., M.O., Y.Y., and T. Saio analyzed the data. N. Iguchi, N. Isozumi, Y.Y., T. Saio, and E.M. wrote the paper. T.K., N.E., M.Y., M.N., S.O., I.O., N.S., M.A., and K.S. helped to analyze and interpret the data, and critically revise the manuscript. K. Sugie, and E.M. conceptualized the study, developed the study design, supervised the authors throughout the study, and provided expertise in manuscript preparation. All authors read and approved the final manuscript.

## Competing interests

EM is a Founder CEO of molmir, Inc..

