## Extended Data for "Zinc finger domains bind low-complexity domain polymers"

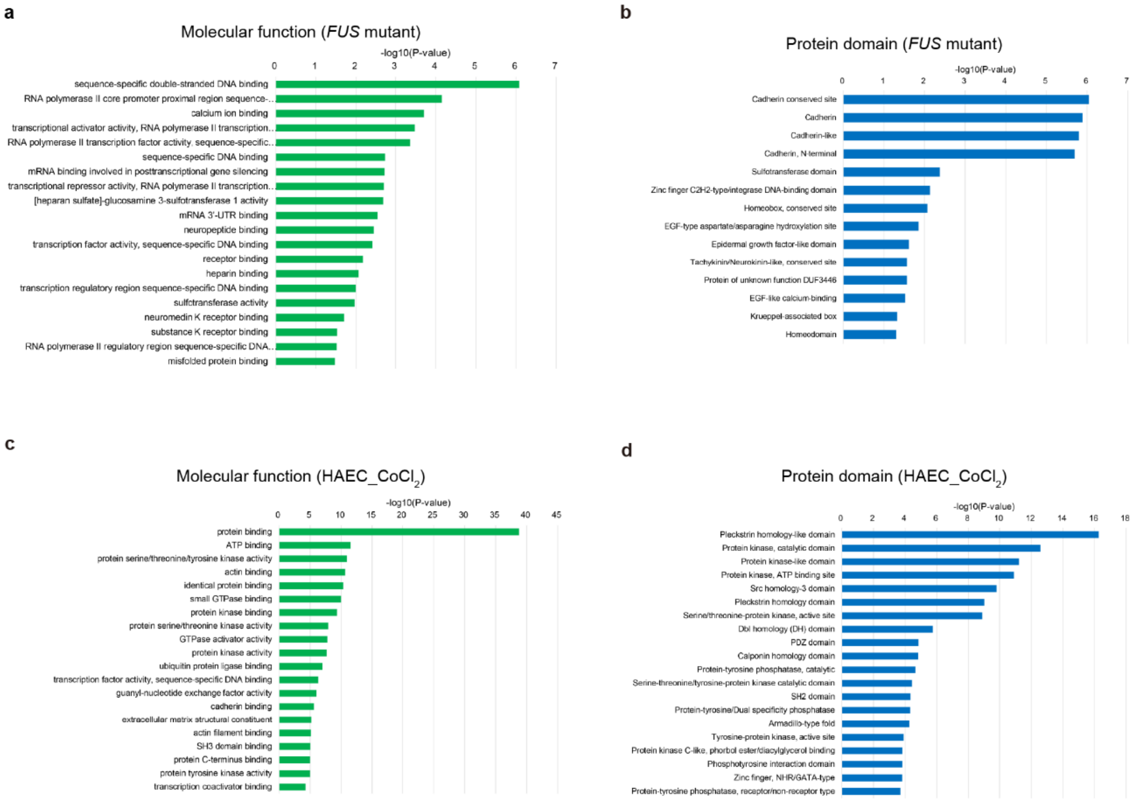


**Extended Data Fig. 1: Gene ontology and protein domain enrichment analyses. a.** GO analysis of molecular function term for *FUS* mutant data. **b.** Protein domain analysis for *FUS* mutant data. **c**. GO analysis of molecular function term for HAEC_$CoCl_2$ data. **d.** Protein domain analysis for HAEC_$CoCl_2$ data.

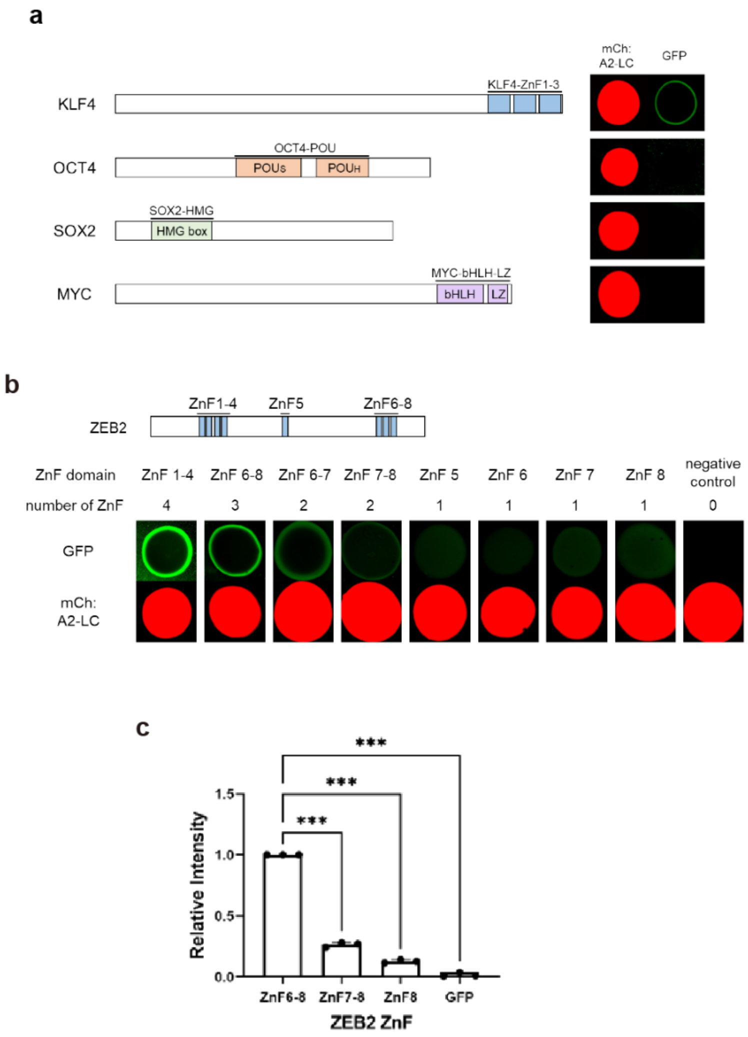


**Extended Data Fig. 2: Hydrogel binding assays. a.** Binding assays of GFP fusion DNA-binding domains to mCh:A2-LC hydrogels. **b.** Binding assays of GFP fusion ZEB2 ZnF to mCh:A2-LC hydrogels. **c.** Quantitative analysis of binding assays (b). Relative intensities of GFP:ZnF6-8, GFP:ZnF7-8, GFP:ZnF8, and GFP are shown in bar charts. Data represent means ± SD ($n$ = 3), analyzed by one-way ANOVA followed by Dunnett's multiple comparison test (***P < 0.001).

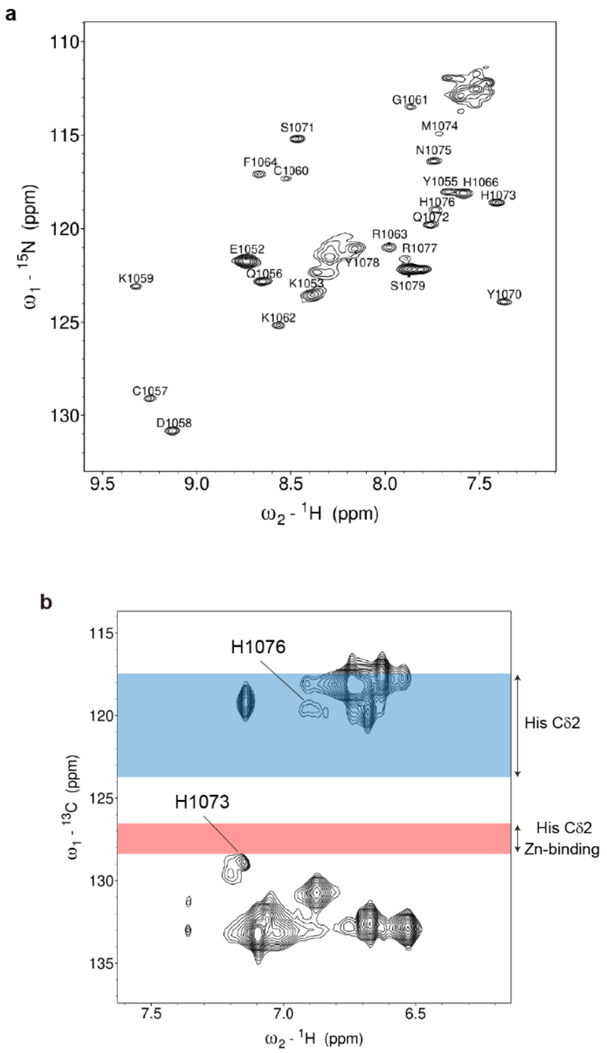


**Extended Data Fig. 3: NMR analyses of ZEB2 ZnF8. a**. Signal assignment of ZnF8. **b**. The chemical shift of H1073 side chain.

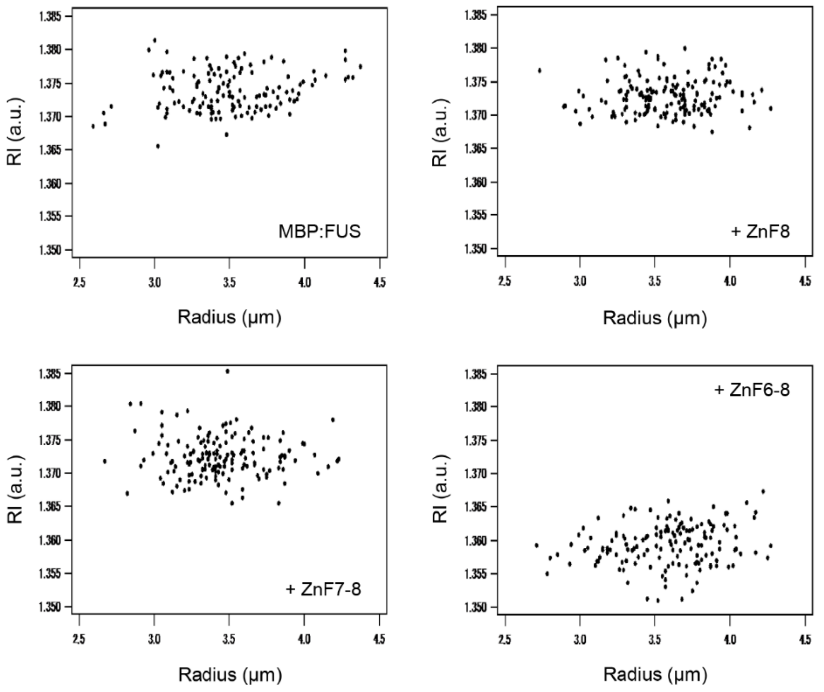


**Extended Data Fig. 4: The size of MBP:FUS droplets in the absence and presence of ZEB2 ZnF.**

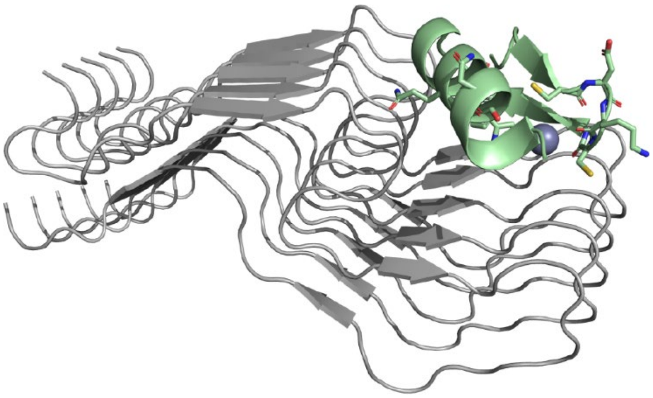

**Extended Data Fig. 5: Binding model of ZnF and LC fibril.** MD simulations of ZEB2 ZnF8 (green) bound to the LC domain of hnRNPA2 (gray) were performed and the resulting model is shown as a ribbon model.
